# The allometry of brain size in mammals

**DOI:** 10.1101/440560

**Authors:** Joseph Robert Burger, Menshian Ashaki George, Claire Leadbetter, Farhin Shaikh

## Abstract

Why some animals have big brains and others do not has intrigued scholars for millennia. Yet, the taxonomic scope of brain size research is limited to a few mammal lineages. Here we present a brain size dataset compiled from the literature for 1552 species with representation from 28 extant taxonomic orders. The brain-body size allometry across all mammals is (*Brain*) = −1.26 (*Body*)^0.75^. This relationship shows strong phylogenetic signal as expected due to shared evolutionary histories. Slopes using median species values for each order, family, and genus, to ensure evolutionary independence, approximate ∼0.75 scaling. Why brain size scales to the ¾ power to body size across mammals is, to our knowledge, unknown. Slopes within taxonomic orders exhibiting smaller size ranges are often shallower than 0.75 and range from 0.24 to 0.81 with a median slope of 0.64. Published brain size data is lacking for the majority of extant mammals (>70% of species) with strong bias in representation from Primates, Carnivores, Perrisodactyla, and Australidelphian marsupials (orders Dasyuromorphia, Diprotodontia, Peramelemorphia). Several orders are particularly underrepresented. For example, brain size data are available for less than 20% of species in each of the following speciose lineages: Soricomorpha, Rodentia, Lagomorpha, Didelphimorphia, and Scandentia. Use of museum collections can decrease the current taxonomic bias in mammal brain size data and tests of hypothesis.

---

Brain size in mammals has received attention from scholars for at least 3,500 years since Aristotle made the conjecture that humans have the largest brain in proportion to our size (Striedter 2005). Since Aristotle, brain size has continued to intrigue observers of the natural world including Darwin (1871), Snell (1891), Dubois (1898), Jerison (1973), Gould 1975), Lande (1979) and many studies in recent decades. The accumulation of years of inquiry shows Aristotle’s conjecture is accurate only when considering allometric scaling effects of body size (Figure 1). When allometry is accounted, *Homo sapiens* has the largest brain size of any mammal, and primates in general have relatively large brains compared to other mammals. However, understanding the evolutionary significance of brain size scaling and its deviations across the diversity of mammals remains a challenge. This is in part because much brain size research has focused on limited taxa with a bias towards understanding the evolution of big brains.

**Fig 1.**
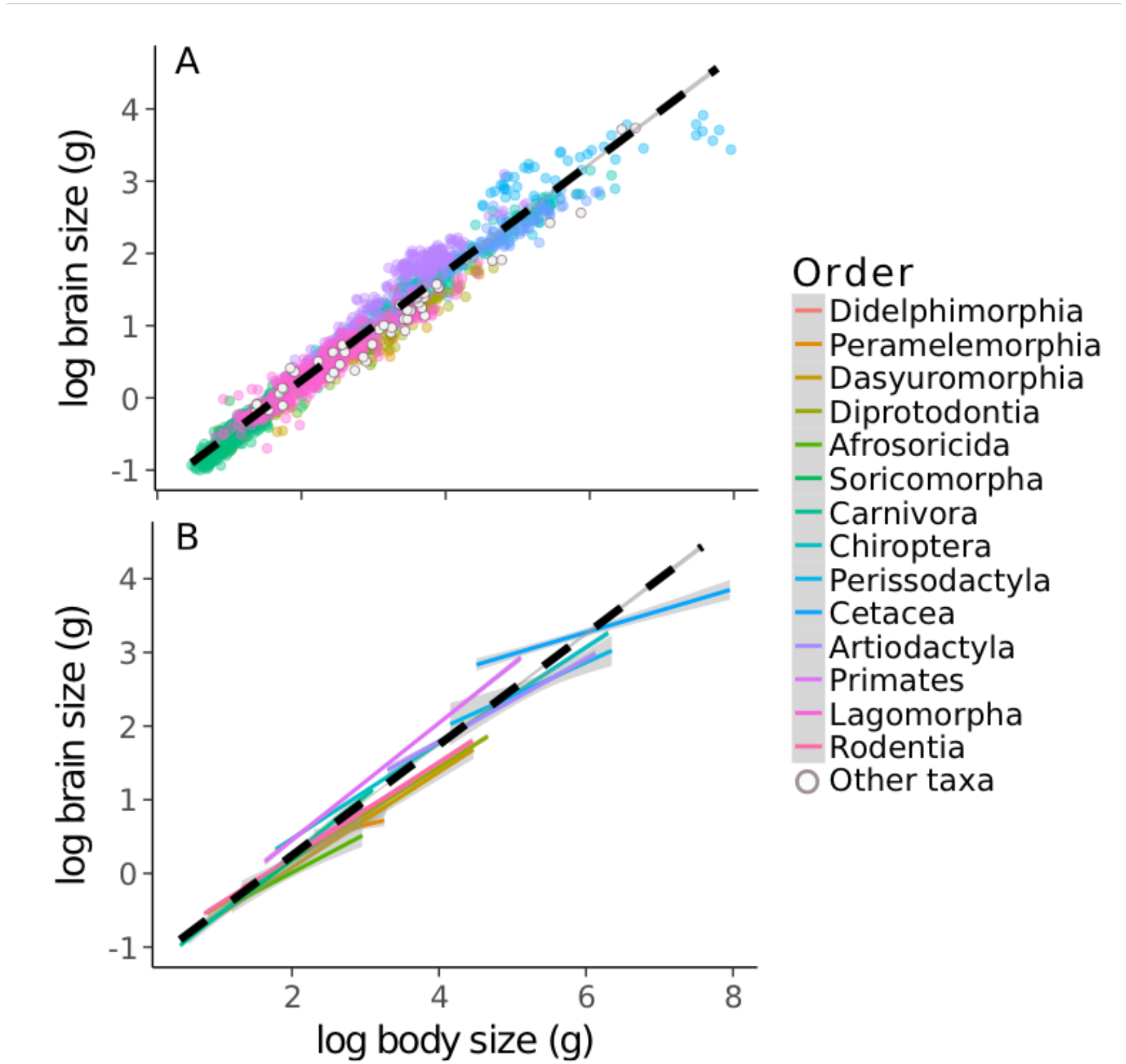
A) The allometry of brain size versus body size for 1552 mammal species. Points are color coded based on orders with >10 species; open circles are species in orders with ≤10 species. B) The allometries by taxonomic groups (>10 species). Grey bands represent the 95% confidence intervals.

Studies of brain size variation in mammals include a number of hypotheses relating large brain size to various social (e.g., Dunbar and Shultz 2007), ecological (e.g., Sol et al. 2008), energetic (e.g., Isler and Van Schaik 2006), life history (e.g., González-Lagos et al. 2010), and behavioral characteristics (e.g., Benson-Amram et al. 2016). However, the taxonomic scope that brain size hypotheses have been evaluated is limited and often shows mixed results. For example, the social brain hypothesis (Dunbar and Shultz 2007) that linked large brain size in primates to the challenges of social living, has been very popular. However, recent analysis with larger datasets, advanced statistical methods, and alternative hypothesis testing suggests that ecology, including diets of dispersed high quality food items, is a better predictor of brain size than sociality among primates (DeCasien et al. 2017; González-Forero and Gardner 2018). There is lack of support for the social brain hypothesis in other diverse taxonomic groups such as marmots (Matějů et al. 2016), carnivores (Benson-Amram et al. 2016), and bats (Pitnick et al. 2006). In contrast, it is well documented that toothed-whales (Odontoceti) have relatively large brain size and complex socio-ecological systems (Marino 1998; Boddy et al. 2012; Montgomery et al. 2013; Fox et al. 2017). It’s interesting to note that the eusocial mole-rat (*Heterocephallus glaber*) has relatively small brains (Kverková et al. 2018). Moreover, Carnivore brain size is related to puzzle-solving abilities and not sociality (Benson-Amram et al. 2016). These and other studies show that understanding the links between brain size and socio-ecological lifestyles in mammals remains a challenge (Dunbar and Shultz 2017).

The use of brain size in comparative studies is certainly not void of contention and Healy and Rowe (2007) and Logan et al. (2018) provide critiques. We suggest that brain size is still a useful biological trait in comparative and macroecological studies for several reasons. First, across species, brain size varies by more than an order of magnitude even after accounting for strong allometric scaling effects of body size (Figure 1, Figure 2). So, this variation cannot simply be due to measurement error and therefore begs a biological explanation. Second, the brain is an energetically expensive organ, appropriating about ∼20% of energy use in humans (or ∼400 kilocalories per day) while only contributing to ∼2% of human body mass (Herculano-Houzel 2011). Total brain size represents the aggregate cognitive costs of all of the brain parts, and thus, variation in brain size is integral to understanding how animals allocate resources to various tradeoffs and life history traits that affect survival and reproduction. Third, brain size in mammals is unambiguous and easily quantified from museum specimens allowing for large sample sizes spanning many taxa which are required for macroecological and comparative studies. Moreover, these allometries from modern mammals are often used in studying fossil mammals (e.g., Finarelli and Flynn 2009; Smaers et al. 2012). So, quantifying the overall and taxon-specific allometries (Pagel and Harvey 1989) are important for inferring basic biology and natural history of modern and extinct mammals.

**Fig 2.**
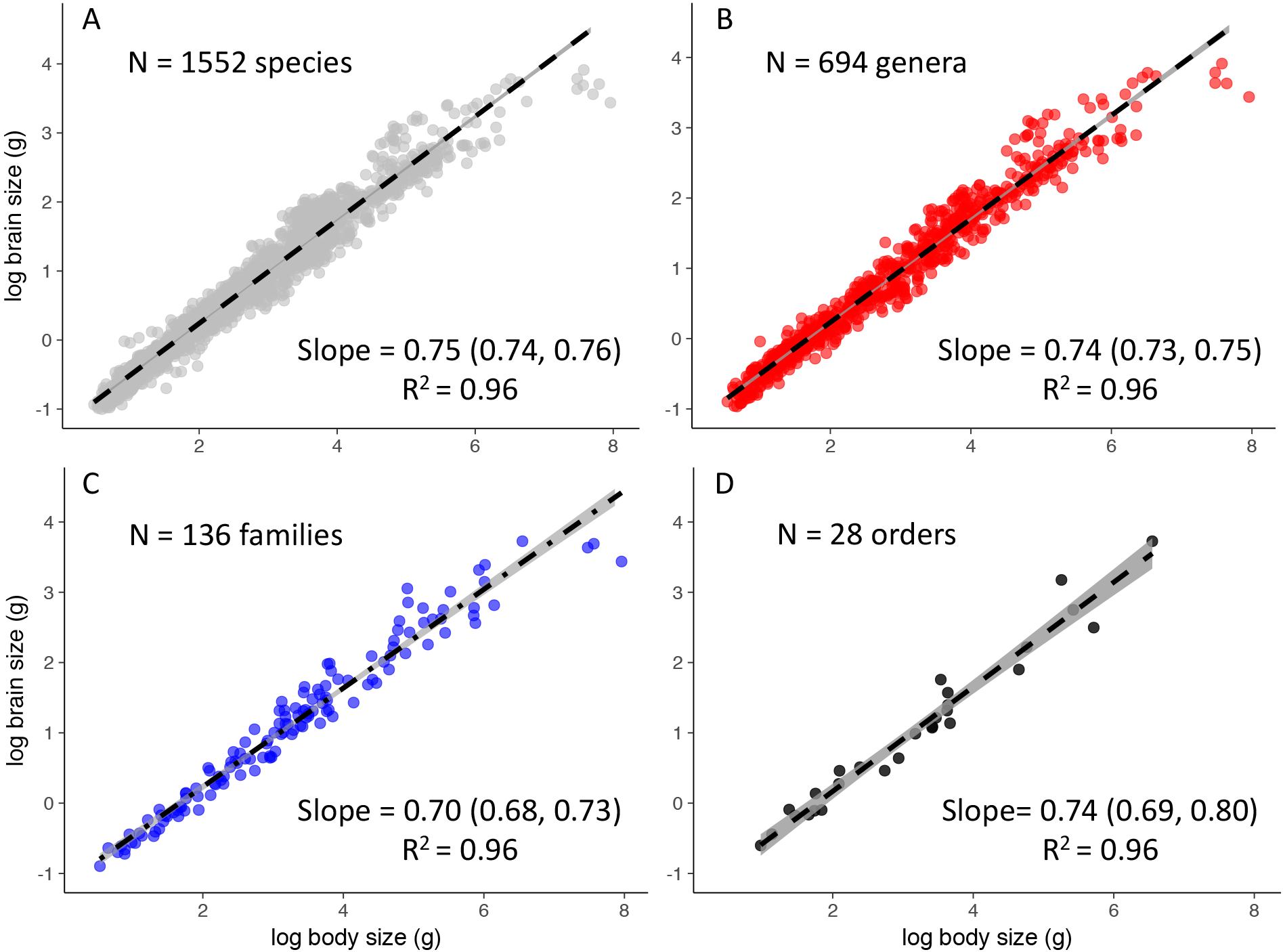
The allometry of brain size to body size across all species (A) and with median values by genera (B), families (C), and orders (D). Grey bands represent the 95% confidence intervals. Note that confidence intervals at all taxonomic scales are statistically indistinguishable from 0.75 with the exception of family-level allometry.

Here we present a brain size dataset compiled from the literature for 1552 species spanning 28 extant taxonomic orders. We analyze the allometry of brain size to body size across all mammals and at multiple evolutionary scales. We summarize the general patterns in relative brain size variation among and within major evolutionary lineages and lifestyle groups. We end by highlighting some questions and gaps in brain size literature offering future research opportunities.

## MATERIALS AND METHODS

### Brain size data

We conducted an extensive search using google scholar and datadryad.org for published brain size datasets and other publications reporting brain size measurements. Data inclusion was based on the following criteria. We used brain size and body size data from the same published source when possible. We referenced body size data when from a different source than brain size. We report sex and sample sizes or ranges (e.g., 1<N<10) of adult animals used in estimating brain size when reported. We used averages for adults of both sexes, and adult female brain and body size for lineages known to exhibit sexual size dimorphism following (Isler and van Schaik 2012). For published datasets, we verified references for accuracy and merged data into a master file standardized by taxonomy in (Wilson and Reeder 2005). When subspecies were reported, we took mean values for species weighted for sample sizes. We used a conversion of 1 gram to 1 cm^3^ when different units were reported following earlier studies (e.g., Isler and van Schaik 2009). The final data set includes brain size and body size estimates for 1552 mammals collated and verified from 54 published references. For each entry, we include taxonomy (Order, Family, Genus, and Latin binomial) mean brain size (g), mean body size (g), brain size residuals from the overall allometry (e.g., *Homo sapiens* compared to all other mammals), and order-specific residuals (e.g, *H. sapiens* compared to other primates (see supplemental materials for dataset).

### Analysis

The scaling of brain size with body size is typically characterized by a power law (Snell 1891; Dubois, E. 1898; Jerison 1973), where (*Brain Size*) = *α* (*Body Size*)^*β*^ where *α* and *β* are constants representing the intercept and slope, respectively. This relationship becomes linear by log transforming both sides of the equation, *log* (*Brain Size*) = *log*(*α*) − *β* × *log* (*Body Size*). Ordinary Least Squares (OLS) Regressions fitted to data allow comparisons of parameter estimates (intercept and slope) across species and taxonomic and lifestyle groups.

We first evaluate the power law scaling of brain size to body size across all species (N = 1552) in the dataset using OLS. We then conducted a Phylogenetic Generalized Least Squares (PGLS) using the mammal supertree from (Bininda-Emonds et al. 2007) and (Fritz et al. 2009). This analysis revealed strong phylogenetic signal as expected because evolutionary histories impose limits to how quickly traits can evolve resulting in non-independence among closely related species (Felsenstein 1985). To address this issue, we first evaluated additional allometries using only the median species by order (N = 28), family (N = 136), and genera (N = 694). This prevents speciose lineages with similar lifestyles, brain and body size (e.g., Myomorpha rodents) from driving the regression analysis. Secondly, we conducted taxon-level allometries following Pagel and Harvey (1988) within orders for those with > 10 species with brain size data. This shows how different taxonomic and ‘lifestyle’ groups compare to each other in allometric parameter estimates (Sibly and Brown 2007). We separate cetaceans and artiodactyl because of distinct functional ‘lifestyles’ despite lack of monophyly in the latter. Finally, we plotted violin plots of brain size residuals from the allometry (observed-expected) for each species in the data to visually show the variation in relative brain size among major taxonomic and lifestyle groups. All analyses were conducted in R using the following packages: ‘ggplot2’, ‘phytools’, and ‘caper’, ‘dplyr’ ‘nlme’ (R code and data will be made available in supplemental materials).

## RESULTS

The allometry of brain size to body size across mammal species in our datasets is (*Brain*) = −1.26 (*Body*)^0.75^ (Figure1). It is important to note that the confidence intervals of the slope (0.742, 0.758) exclude linear scaling (slope = 1) and two-thirds scaling. Phylogenetic analysis showed strong signal: Pagel’s λ = 0.935 (Pagel 1999) and allometric parameters intercept = −0.83 and slope = 0.57.

Allometries at coarser taxonomic scales using median species at the order, family, and genera levels, however, approximate 3/4 scaling (Figure 2). Residual deviations about this allometry vary by taxonomic and lifestyle group (Figure 3). Allometric slopes by orders are generally shallower and range from 0.24 in the Australidelphia marsupial order Peramelemorphia (Bandicoots and bilberries) to 0.81 in Chiroptera (bats) with a median value of 0.64 (Table 1). There is also variation in the elevation (intercepts) of the slopes by taxonomic groups. Primates and bats have steeper allometries compared to other orders; primates are significantly above the line for all mammals (Table 1). Lagomorpha and Soricomorpha also have similar slopes to the overall allometry but with lower intercepts.

**Fig 3.**
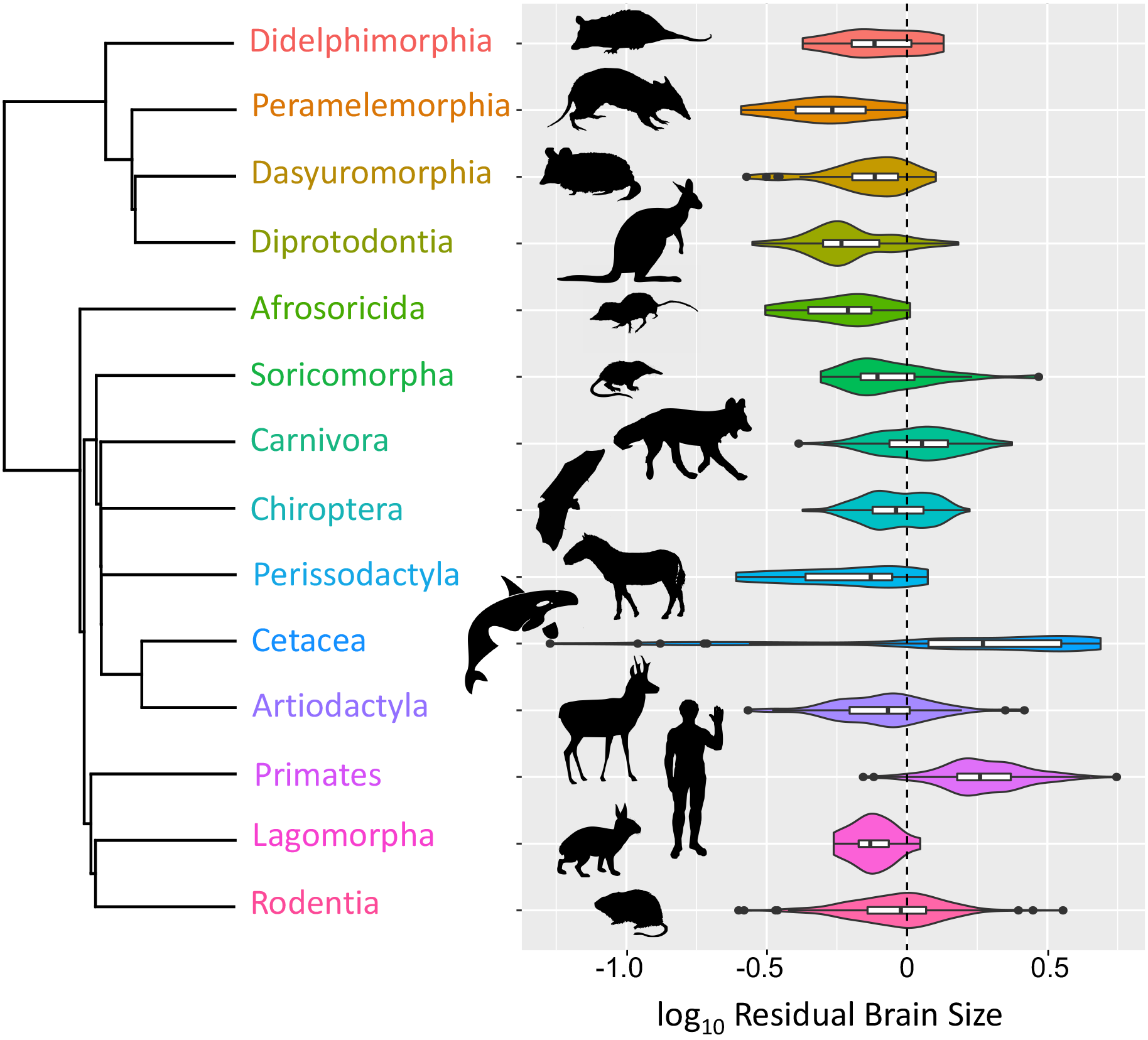
The phylogenetic distribution and violin plots of residual brain sizes by taxonomic order. Plots show mirror curves of kernel density estimation of the data. Inserted box and whisker plots show range (whiskers), interquartile ranges (white boxes), median values (horizontal black lines) of the data. Circles in plot tails indicate outliers. Vertical dashed line at 0 corresponds with dashed lines in figures 1. Taxonomy based on Wilson and Reader (2005) and mammal supertree from Bininda-Emonds et al. (2007) and Fritz et al. (2009).

**Table 1.** Allometries for log10 brain size versus log10 body size across mammals and orders with >10 species for which we have brain size data. CI = confidence interval.

| **Order** | **R<sup>2</sup>** | **N = species** | **Slope [95% CI]** | **Intercept [95% CI]** |
| --- | --- | --- | --- | --- |
| Afrosoricida | 0.85 | 13 | 0.54 [0.38, 0.699] | -1.07 [-1.4, -0.743] |
| Artiodactyla | 0.89 | 95 | 0.56 [0.518, 0.599] | -0.44 [-0.637, -0.246] |
| Carnivora | 0.96 | 199 | 0.65 [0.632, 0.669] | -0.83 [-0.908, -0.762] |
| Cetacea | 0.74 | 43 | 0.29 [0.238, 0.35] | 1.51 [1.19, 1.83] |
| Chiroptera | 0.93 | 309 | 0.81 [0.786, 0.835] | -1.37 [-1.41, -1.34] |
| Dasyuromorphia | 0.95 | 50 | 0.64 [0.597, 0.687] | -1.21 [-1.3, -1.12] |
| Didelphimorphia | 0.93 | 16 | 0.56 [0.472, 0.652] | -0.94 [-1.15, -0.792] |
| Diprotodontia | 0.95 | 112 | 0.63 [0.604, 0.658] | -1.08 [-1.17, 0.99] |
| Lagomorpha | 0.96 | 15 | 0.75 [0.742, 0.759] | -1.26 [-1.29, -1.24] |
| Peramelemorphia | 0.43 | 14 | 0.24 [0.056, 0.423] | -0.06 [-0.594, 0.466] |
| Perissodactyla | 0.44 | 11 | 0.35 [0.0286, 0.67] | 0.74 [-1.06, 2.54] |
| Primates | 0.92 | 248 | 0.79 [0.764, 0.825] | -1.14 [-1.24, -1.03] |
| Rodentia | 0.92 | 351 | 0.64 [0.622, 0.663] | -1.06 [-1.11, -1.02] |
| Soricomorpha | 0.88 | 28 | 0.75 [0.634, 0.864] | -1.32 [-1.47, -1.16] |
| **All mammals** | **0.96** | **1552** | **0.75 [0.742, 0.758]** | **-1.26 [-1.28, -1.24]** |

Published brain size data is lacking for the majority of extant mammals (>70% of species; Table 2), with the best representation (>65% of species) in the following orders: Primate, Carnivora, Perrisodactyla and the Australidelphian marsupials (orders Dasyuromorphia, Diprotodontia, Peramelemorphia). Several orders are underrepresented in the brain size data, including Soricomorpha, Rodentia, Lagomorpha, Didelphimorphia, Scandentia, all having less than 20% of species represented in the brain size data.

## DISCUSSION

### Brain size allometry and its deviations

Across taxa, the three-fourths scaling reveals economies of scale where brain size increases sublinear to body size. There is strong phylogenetic signal in the residual deviations in Figure 1 and PGLS analysis reveals a shallower slope. This is evident in the variation among different lifestyles among taxonomic groups, possibly reflecting development, physiological, and ecological constraints on morphology and anatomical design. For example, in addition to Primates, Carnivora, tree-shrews of the order Scandentia, and the Odontocete cetaceans also show large brain size as previously reported. In contrast, manatees, Australian marsupials, and Perissodactyls have relatively small brains. Rodents and Artiodactyls have medium sized brains although these orders show
the greatest variation in brain size. Cetaceans have the largest variation in residual values of any order, with baleen whales having relatively small brains and toothed whales have large brains (Boddy et al. 2012; Fox et al. 2017; Figure 3).

We conducted additional scaling analyses using only median values at the order and family levels representing to ensure independent evolutionary units. These allometries show consistant ∼¾ scaling when removing potential biases of highly speciose lineages driving the allometry. This differs from some studies that report ∼2/3 scaling (e.g., Dubois, E. 1898; Jerison 1973; Sol et al. 2008) but consistent with other studies reporting ∼¾ scaling (Isler and van Schaik 2009; Boddy et al. 2012; Stankowich and Romero 2017). A recent large comparative analysis reports ∼0.5 scaling using phylogenetic analysis (Tsuboi et al. 2018),which is similar to our 0.57 scaling using PGLS. This is likely because slopes within taxonomic orders are typically shallower than median values among orders (Table 1) as noted earlier (Pagel and Harvey 1988). Notable exceptions, however, are bats and primates which show steeper slopes (Table 1). Lagomorphs and Soricomorpha also have slopes ∼0.75 and intercepts below the allometry for all mammals. Previous authors (e.g., Lande 1979; Pagel and Harvey 1988; Smaers et al. 2012; Tsuboi et al. 2018) have discussed the various selection pressures that act on both brain size as well as body size providing different scenarios of correlated trait evolution. With the emergence of new datasets and statistical approaches, opportunities abound to understanding the role of selection on both brain and body size in producing relative brain size across phylogenetic scales (Smaers et al. 2012; Tsuboi et al. 2018).

### Filling the gaps in brain size data

We have incomplete data available on brain size with roughly 70% of mammal diversity missing, with clear taxonomic bias in brain size data in the literature (Table 2). This dataset is biased with best representation (>65% of species) from Australidelphia marsupials (Dasyuromorphia, Diprotodontia, Peramelemorphia), primates (66.22% of species), carnivores (70% of species), and perrisodactyla (65%). Brain size is typically quantified from the endocranial volume of skulls by plugging holes and filling the cavity with seeds, buckshot, or glass beads and then decanted into a graduated cylinder to measure volume (Gittleman 1986; Iwaniuk and Nelson 2002). Brain size is occasionally measured as wet mass for freshly killed animals (e.g., Boddy et al. 2012), and comparisons between techniques using wet mass and endocranial volume from skulls have been validated ((Iwaniuk and Nelson 2002; Logan and Clutton-Brock 2013). Museum specimens can play an important role in reducing the bias in brain size data in the literature.

### Why ¾ scaling

Empirical scaling patterns in mammals have been central to recent advancements towards ‘universal scaling laws’ that seek to integrate form and function across levels of biological processes from physiology morphology and behavior to life history, populations, and ecosystems (Sibly et al. 2012). Understanding the evolutionary significance of brain size scaling has much potential to unite the common currencies of energy and information to understand the dual processes governing complex biological systems. The scaling of brain size with body size to the three-fourths power mirrors the scaling of metabolic rate with body size that is central to the metabolic theory of ecology (Brown et al. 2004). However, the link between brain size and metabolic scaling is not so clear. Several studies have tested the correlation between residual deviations in brain size and metabolic rate when controlling for body size. These results generally support a positive relationship between relative brain size and metabolic rate (Martin 1981; Armstrong 1983), however the correlation is weak (e.g., Isler and van Schaik 2006) and variable at different taxonomic scales and sensitive to phylogenetic analyses (Sobrero et al. 2011). It should be noted taht these are only two parameters in the overall energy budget. Understanding how brain size fits into a complete energetic framework that captures tradeoffs with other major energy demands including other expensive tissues (e.g., Aiello and Wheeler 1995; Navarrete et al. 2011) and life history traits (e.g., Isler and van Schaik 2012) that contribute to survival and reproduction is still needed.

The allometry of brain size is unique among organs which typically scale with steeper slopes or isometrically with body size (Peters and Peters 1986). Organs such as the stomach, heart, lungs, and liver
are highly vascularized and feature fractal-like resource distribution networks that produce ¾ scaling (West et al. 1997; Banavar et al. 2010). While the size of these organs scale linearly with body size, their metabolic rates scale to the ∼¾ power, consistent with metabolic scaling theory. In contrast, the mammalian brain scales sublinearly with body size. It is possible that this is a result of increased modularity and folding that allows for the observed economies of scale in energy use with increasing size. There is evidence from FMRI scans from few mammals that the basal metabolic rate of the brain scales steeper and close to linear (∼5/6) with brain size (Karbowski 2007). Energy use scales linearly with number of neurons which is highly correlated with brain size (Herculano-Houzel 2011). So, a possible unique characteristic of the brain, is that brain metabolism scales linearly with brain size while the brain scales sublinear to body size. This would result in the scaling of total energy allocation to the brain approximating ¾ as with other organs showing isometry with body size. Understanding the scaling of size and metabolic costs of the brain and other organs across body sizes and evolutionary lineages is much needed to further understand the significance of sublinear scaling of brain size in mammals.

## ACKNOWLEDGEMENTS

MJ Hamilton, JH Brown, PR Stephens, AD Davidson, TS Fristoe, JM Grady, FA Smith, and AH Hurlbert provided helpful discussions. We thank John Gittleman, Simon Reader, and Karin Isler for providing versions of published datasets that we cross-checked in compiling our dataset. We acknowledge funding from an American Society of Mammalogists Shadle Fellowship, The McNair Scholars Program, and a UNC Carolina Postdoctoral Fellowship for Faculty Diversity awarded to JRB.

## Supplemental materials

1. Brain size data (email corresponding author for dataset)
2. Brain size references

## Appendix 1

Percent of orders with brain size data. Taxonomy based on Wilson and Reader (2005).

| Order | # of species | Species with brain data | % of order with brain data |
| --- | --- | --- | --- |
| Rodentia | 2277 | 351 | 15.42 |
| Chiroptera | 1116 | 309 | 27.69 |
| Soricomorpha | 428 | 28 | 6.54 |
| Primates | 376 | 248 | 65.96 |
| Carnivora | 286 | 199 | 69.58 |
| Artiodactyla | 240 | 95 | 39.58 |
| Diprotodontia | 143 | 112 | 78.32 |
| Lagomorpha | 92 | 15 | 16.30 |
| Didelphimorphia | 87 | 16 | 18.39 |
| Cetacea | 84 | 43 | 51.19 |
| Dasyuromorphia | 71 | 50 | 70.42 |
| Afrosoricida | 51 | 13 | 25.49 |
| Erinaceomorpha | 24 | 7 | 29.17 |
| Peramelemorphia | 21 | 14 | 66.67 |
| Cingulata | 21 | 7 | 33.33 |
| Scandentia | 20 | 4 | 20.00 |
| Perissodactyla | 17 | 11 | 64.71 |
| Macroscelidea | 15 | 5 | 33.33 |
| Pilosa | 10 | 7 | 70.00 |
| Pholidota | 8 | 3 | 37.50 |
| Paucituberculata | 6 | 2 | 33.33 |
| Monotremata | 5 | 3 | 60.00 |
| Sirenia | 5 | 2 | 40.00 |
| Hyracoidea | 4 | 3 | 75.00 |
| Proboscidea | 3 | 2 | 66.67 |
| Notoryctemorphia | 2 | 1 | 50.00 |
| Dermoptera | 2 | 0 | 0.00 |
| Microbiotheria | 1 | 1 | 100.00 |
| Tubulidentata | 1 | 1 | 100.00 |
| **TOTAL** | **5416** | **1552** | **28.67** |

## Literature Cited

Aiello, L. C., and P. Wheeler. 1995. The expensive-tissue hypothesis: the brain and the digestive system in human and primate evolution. Current Anthropology 36:199–221.

Armstrong, E. 1983. Relative brain size and metabolism in mammals. Science 220:1302–1304.

Banavar, J. R. et al. 2010. A general basis for quarter-power scaling in animals. Proceedings of the National Academy of Sciences 107:15816–15820.

Benson-Amram, S., B. Dantzer, G. Stricker, E. M. Swanson, and K. E. Holekamp. 2016. Brain size predicts problem-solving ability in mammalian carnivores. Proceedings of the National Academy of Sciences 113:2532–2537.

Bininda-Emonds, O. R. et al. 2007. The delayed rise of present-day mammals. Nature 446:507.

Boddy, A. M., M. R. Mcgowen, C. C. Sherwood, L. I. Grossman, M. Goodman, and D. E. Wildman. 2012. Comparative analysis of encephalization in mammals reveals relaxed constraints on anthropoid primate and cetacean brain scaling. Journal of Evolutionary Biology 25:981–994.

Brown, J. H., J. F. Gillooly, A. P. Allen, V. M. Savage, and G. B. West. 2004. Toward a metabolic theory of ecology. Ecology 85:1771–1789.

Darwin, C. 1871. The descent of man. The Great Books of the Western World 49:320.

Decasien, A. R., S. A. Williams, and J. P. Higham. 2017. Primate brain size is predicted by diet but not sociality. Nature Ecology & Evolution 1:0112.

Dubois, E. 1898. Ueber die Abhängigkeit des Hirngewichtes von der Körpergrösse bei den Säugethieren. Archiv für Anthropologie 25:1–28.

Dunbar, R. I., and S. Shultz. 2007. Evolution in the social brain. Science 317:1344–1347.

Felsenstein, J. 1985. Phylogenies and the comparative method. The American Naturalist 125:1–15.

Finarelli, J. A., and J. J. Flynn. 2009. Brain-size evolution and sociality in Carnivora. Proceedings of the National Academy of Sciences 106:9345–9349.

Fox, K. C., M. Muthukrishna, and S. Shultz. 2017. The social and cultural roots of whale and dolphin brains. Nature Ecology & Evolution 1:1699.

Fritz, S. A., O. R. Bininda-Emonds, and A. Purvis. 2009. Geographical variation in predictors of mammalian extinction risk: big is bad, but only in the tropics. Ecology letters 12:538–549.

Gittleman, J. L. 1986. Carnivore brain size, behavioral ecology, and phylogeny. Journal of Mammalogy 67:23–36.

González-Forero, M., and A. Gardner. 2018. Inference of ecological and social drivers of human brain-size evolution. Nature 557:554.

González-Lagos, C., D. Sol, and S. M. Reader. 2010. Large-brained mammals live longer. Journal of Evolutionary Biology 23:1064–1074.

Gould, S. J. 1975. Allometry in primates, with emphasis on scaling and the evolution of the brain. Contributions to Primatology 5:244–292.

Healy, S. D., and C. Rowe. 2007. A critique of comparative studies of brain size. Proceedings of the Royal Society of London B: Biological Sciences 274:453–464.

Herculano-Houzel, S. 2011. Scaling of brain metabolism with a fixed energy budget per neuron: implications for neuronal activity, plasticity and evolution. PLoS One 6:e17514.

Isler, K., and C. P. Van Schaik. 2009. The expensive brain: a framework for explaining evolutionary changes in brain size. Journal of Human Evolution 57:392–400.

Isler, K., and C. P. Van Schaik. 2012. Allomaternal care, life history and brain size evolution in mammals. Journal of Human Evolution 63:52–63.

Isler, K., and C. P. Van Schaik. 2006. Metabolic costs of brain size evolution. Biology Letters 2:557–560.

Iwaniuk, A. N., and J. E. Nelson. 2002. Can endocranial volume be used as an estimate of brain size in birds? Canadian Journal of Zoology 80:16–23.

Jerison, H. J. 1973. Evolution of the brain and intelligence Academic Press.

Karbowski, J. 2007. Global and regional brain metabolic scaling and its functional consequences. BMC Biology 5:18.

Kverková, K. et al. 2018. Sociality does not drive the evolution of large brains in eusocial African mole-rats. Scientific Reports 8:9203.

Lande, R. 1979. Quantitative genetic analysis of multivariate evolution, applied to brain: body size allometry. Evolution 33:402–416.

Logan, C. J. et al. 2018. Beyond Brain Size: uncovering the neural correlates of behavioral and cognitive specialization. Comparative Cognition & Behavior Reviews 13.

Logan, C. J., and T. H. Clutton-Brock. 2013. Validating methods for estimating endocranial volume in individual red deer (Cervus elaphus). Behavioural Processes 92:143–146.

Marino, L. 1998. A comparison of encephalization between odontocete cetaceans and anthropoid primates. Brain, Behavior and Evolution 51:230–238.

Martin, R. D. 1981. Relative brain size and basal metabolic rate in terrestrial vertebrates. Nature 293:57–60.

Matějů, J., L. Kratochvíl, Z. Pavelková, V. P. Řičánková, V. Vohralík, and P. Němec. 2016. Absolute, not relative brain size correlates with sociality in ground squirrels. Proc. R. Soc. B 283:20152725.

Montgomery, S. H., J. H. Geisler, M. R. Mcgowen, C. Fox, L. Marino, and J. Gatesy. 2013. The evolutionary history of cetacean brain and body size. Evolution 67:3339–3353.

Navarrete, A., C. P. Van Schaik, and K. Isler. 2011. Energetics and the evolution of human brain size. Nature 480:91.

Pagel, M. 1999. Inferring the historical patterns of biological evolution. Nature 401:877.

Pagel, M. D., and P. H. Harvey. 1988. The taxon-level problem in the evolution of mammalian brain size: facts and artifacts. The American Naturalist 132:344–359.

Pagel, M. D., and P. H. Harvey. 1989. Taxonomic differences in the scaling of brain on body weight among mammals. Science 244:1589–1593.

Peters, R. H., and R. H. Peters. 1986. The ecological implications of body size. Cambridge University Press.

Pitnick, S., K. E. Jones, and G. S. Wilkinson. 2006. Mating system and brain size in bats. Proceedings of the Royal Society of London B: Biological Sciences 273:719–724.

Sibly, R. M., and J. H. Brown. 2007. Effects of body size and lifestyle on evolution of mammal life histories. Proceedings of the National Academy of Sciences 104:17707–17712.

Sibly, R. M., J. H. Brown, and A. Kodric-Brown. 2012. Metabolic ecology: a scaling approach. John Wiley & Sons.

Smaers, J. B., D. K. Dechmann, A. Goswami, C. Soligo, and K. Safi. 2012. Comparative analyses of evolutionary rates reveal different pathways to encephalization in bats, carnivorans, and primates. Proceedings of the National Academy of Sciences:201212181.

Snell O. 1891. Die Abhängigkeit des Hirnge-wichts von dem Körpergewicht und den geistigen Fähigkeiten. Arch Psychiatr Ner-venkr 23: 436–446.

Sobrero, R., L. J. May-Collado, I. Agnarsson, and C. E. HernÁndez. 2011. Expensive brains:“brainy” rodents have higher metabolic rate. Frontiers in Evolutionary Neuroscience 3:2.

Sol, D., S. Bacher, S. M. Reader, and L. Lefebvre. 2008. Brain size predicts the success of mammal species introduced into novel environments. The American Naturalist 172:S63–S71.

Stankowich, T., and A. N. Romero. 2017. The correlated evolution of antipredator defences and brain size in mammals. Proc. R. Soc. B 284:20161857.

Striedter, G. F. 2005. Principles of Brain Evolution (Sinauer, Sunderland, MA).

Tsuboi, M. et al. 2018. Breakdown of brain-body allometry and the encephalization of birds and mammals. Nature Ecology & Evolution 2:1492–1500.

West, G. B., J. H. Brown, and B. J. Enquist. 1997. A general model for the origin of allometric scaling laws in biology. Science 276:122–126.

Wilson, D. E., and D. M. Reeder. 2005. Mammal species of the world: a taxonomic and geographic reference. JHU Press.

